# High-fat diet feeding triggers a regenerative response in the adult zebrafish brain

**DOI:** 10.1101/2022.11.22.517521

**Authors:** Yagmur Azbazdar, Yusuf Kaan Poyraz, Ozgun Ozalp, Dilek Nazli, Dogac Ipekgil, Gokhan Cucun, Gunes Ozhan

**Author notes:** These authors contributed equally to this work (Yagmur Azbazdar, Yusuf Kaan Poyraz). These authors contributed equally to this work (Ozgun Ozalp, Dilek Nazli). Correspondence: Gunes Ozhan.

## Abstract

Non-alcoholic fatty liver disease (NAFLD) includes a range of liver conditions ranging from excess fat accumulation to liver failure. NAFLD is strongly associated with high-fat diet (HFD) consumption that constitutes a metabolic risk factor. While HFD has been elucidated concerning its several systemic effects, there is little information about its influence on the brain at the molecular level. Here, by using a high-fat diet (HFD)-feeding of adult zebrafish, we first reveal that excess fat uptake results in weight gain and fatty liver. Prolonged exposure to HFD induces a significant increase in the expression of pro-inflammation, apoptosis, and proliferation markers in the liver and brain tissues. Immunofluorescence analyses of the brain tissues disclose stimulation of apoptosis and widespread activation of glial cell response. Moreover, glial activation is accompanied by an initial decrease in the number of neurons and their subsequent replacement in the olfactory bulb and the telencephalon. Long-term consumption of HFD causes activation of Wnt/β-catenin signaling in the brain tissues. Finally, fish fed an HFD induces anxiety, and aggressiveness and increases locomotor activity. Thus, HFD feeding leads to a non-traumatic brain injury and stimulates a regenerative response. The activation mechanisms of a regeneration response in the brain can be exploited to fight obesity and recover from non-traumatic injuries.

## Introduction

Non-alcoholic fatty liver disease (NAFLD) is a common disorder and is defined as a group of conditions caused by excessive fat accumulation in the liver. NAFLD, characterized by an increase in intrahepatic lipid content, has frequently been associated with a disproportionate amount of dietary fat intake and obesity (Fabbrini et al., 2010). High-fat diet (HFD)-associated obesity is very common in patients with NAFLD and has been exploited to generate animal models of NAFLD (Lian et al., 2020). A growing body of evidence suggests that NAFLD does not remain restricted to the liver, but acts as an early mediator of systemic diseases that affect extrahepatic organs (Byrne and Targher, 2015; Reccia et al., 2017). NAFLD has been reported to adversely affect various organs including those of the gastrointestinal system, the kidney, the heart, and the brain (VanWagner and Rinella, 2016; Vargas and Vásquez, 2017; Li et al., 2020; Rohr et al., 2020; Picolo et al., 2021).

Numerous cross-sectional studies related to increased body mass index, weight gain, and obesity in adults aged 18-65 years have revealed that these phenomena can be related to deficits in certain cognitive functions including psychomotor performance and speed, visual construction, verbal memory, decision making and executive function (Cournot et al., 2006; Fergenbaum et al., 2009; Smith et al., 2011; Fagundo et al., 2012; Prickett et al., 2015). Obesity has also been linked with structural and metabolic alterations in the brain that lead to impairment of cognitive functions (Prickett et al., 2015; Livingston et al., 2020; Sui and Pasco, 2020). These alterations span a range of potential mechanisms including compromised cerebral metabolism, elevated leptin levels, increased inflammation, and neuronal injury (Volkow et al., 2009; Mueller et al., 2012; O’Brien et al., 2017; Yin et al., 2018; Zatterale et al., 2020). The relationship between overfeeding and obesity, and the brain has been investigated in animal models of a high-fat diet (HFD) (Hariri and Thibault, 2010; Zang et al., 2018; de Moura e Dias et al., 2021). Owing to the similarity of the transcriptional, metabolic, and behavioral responses to those of mammals, zebrafish has been evaluated as a valuable model to study the pathological influence of HFD in humans (Vargas and Vásquez, 2017; Carnovali et al., 2018; Meguro et al., 2019; Ka et al., 2020; Picolo et al., 2021). Accordingly, lipid accumulation in the liver has been accompanied by an increase in the expression of pro-inflammation-related genes in the zebrafish and mammalian models fed an HFD. Mouse models of HFD have revealed cognitive, microstructural, neurochemical, and metabolic alterations in the brain (Winocur et al., 2005; Lee et al., 2018; Nakandakari et al., 2019; Zhao et al., 2019; Chiazza et al., 2021; Guadilla et al., 2021; Leyh et al., 2021). However, a comprehensive molecular analysis of the influence of HFD on the brain regions concerning the activation of cellular processes such as proliferation, cell death, and signaling pathways has not been conducted so far.

Owing to its key roles in the development and growth of tissues and organs involved in the regulation of metabolism, Wnt signaling, in particular, the β-catenin pathway (also termed the canonical Wnt pathway) has been associated with a range of metabolic diseases including obesity and NAFLD (Karabicici et al., 2021). Moreover, Wnt/β-catenin signaling plays key roles in the activation of tissue repair mechanisms in a broad range of regenerating tissues and organs studied to date (Whyte et al., 2012; Ozhan and Weidinger, 2014; Ozhan and Weidinger, 2015). Parts of the central nervous system (CNS) including the retina, optic tectum, and spinal cord have been deciphered to activate Wnt signaling to mediate proliferation and differentiation of stem/progenitor cells in response to injury in the adult zebrafish (Meyers et al., 2012; Briona et al., 2015; Shi et al., 2015; Wehner et al., 2017; Shimizu et al., 2018). Our previous study has further supported the role of Wnt/β-catenin signaling in the adult zebrafish telencephalon where the pathway becomes activated at a very early stage of regeneration, i.e. only 20 hours after injury, and controls a large pool of genes involved in p53, apoptosis, MAPK, mTOR and FoxO pathways (Demirci et al., 2020). Nevertheless, whether HFD feeding can stimulate an injury-induced regenerative response in the brain and activate Wnt/β-catenin signaling remains to be elucidated.

To address the influence of HFD feeding on the brain, we exploited the adult zebrafish that has a robust capacity to regenerate its CNS due to its constitutively active neurogenic zones of stem/progenitor cells (Kaslin et al., 2008; Kizil et al., 2012). Zebrafish CNS regenerates via proliferation and differentiation of the radial glial cells (RGCs) and neuroepithelial-like progenitor cells (Grandel et al., 2006; Kaslin et al., 2008; Kizil et al., 2012; Lindsey et al., 2017; Zambusi and Ninkovic, 2020). This exclusive ability of zebrafish to regenerate their brain provides a platform that enables direct observation of the molecular effects of HFD on the brain in a highly regenerative context. Initially, we have validated our zebrafish model of HFD in the liver by increased fat deposition and activation of genes related to inflammation, proliferation, and apoptosis. Our results show that the consequences of fatty liver are not limited to the liver but also visible in the whole brain tissue as an increase in the expression of genes related to inflammation, apoptosis, proliferation, and neurogenesis. The HFD-induced alterations are detectable in different regions of the brain, i.e. the olfactory bulb and the telencephalon, as activation of apoptosis, glial cell response, and proliferation. While the number of neurons decreases in response to short-term HFD intake, the lost neurons are replaced by the newly forming ones after prolonged exposure to HFD. Interestingly, HFD-induced fatty liver results in the activation of Wnt/β-catenin signaling in different regions of the brain, most prominently after long-term HFD intake. Finally, HFD feeding for short- or long-term causes an increase in the anxiety, aggressiveness, and locomotor activity of the fish. Overall, HFD feeding initiates a regenerative response in the brain.

## Materials and Methods

### Zebrafish husbandry

Zebrafish were supplied by Izmir Biomedicine and Genome Center (IBG) Zebrafish Core Facility and were maintained in a 14-hour light/10-hour dark cycle at 28.5 °C, following the guidelines of the IBG Animal Care and Use Committee. Only male zebrafish were used in the experiments to minimize the effect of female reproductive cycles and hormone fluctuations. All animal experiments were performed according to the European Union Directive 2010/63/EU on the protection of animals used for scientific purposes and approved by the Animal Experiments Local Ethics Committee of Izmir Biomedicine and Genome Center (IBG-AELEC).

### Zebrafìsh feeding and experimental design

1-year-old male Golden zebrafish were synchronized by feeding with 5 mg artemia for 1 month before the experiment. The control group was fed a normal diet of 5 mg artemia for two months and a high-fat diet (HFD) group was fed 5 mg artemia and 30 mg egg yolks from chicken (59% fat, 32% proteins, 2% carbohydrates; Sigma-Aldrich, MO, USA) for one or two months (Landgraf et al., 2017). The fish were fed twice a day and ensured that the meals were consumed in 1-2 minutes. Both control and HFD groups contained eight zebrafish. To identify the influence of HFD feeding on the brain at the molecular level, we fed zebrafish an HFD for up to two months. The experimental workflow included the generation of an HFD model, tissue preparation for immunohistochemical/ immunofluorescence analysis of the liver/brain, and dissection of both liver and brain for gene expression analysis by qPCR (Figure 1).

**Figure 1.**
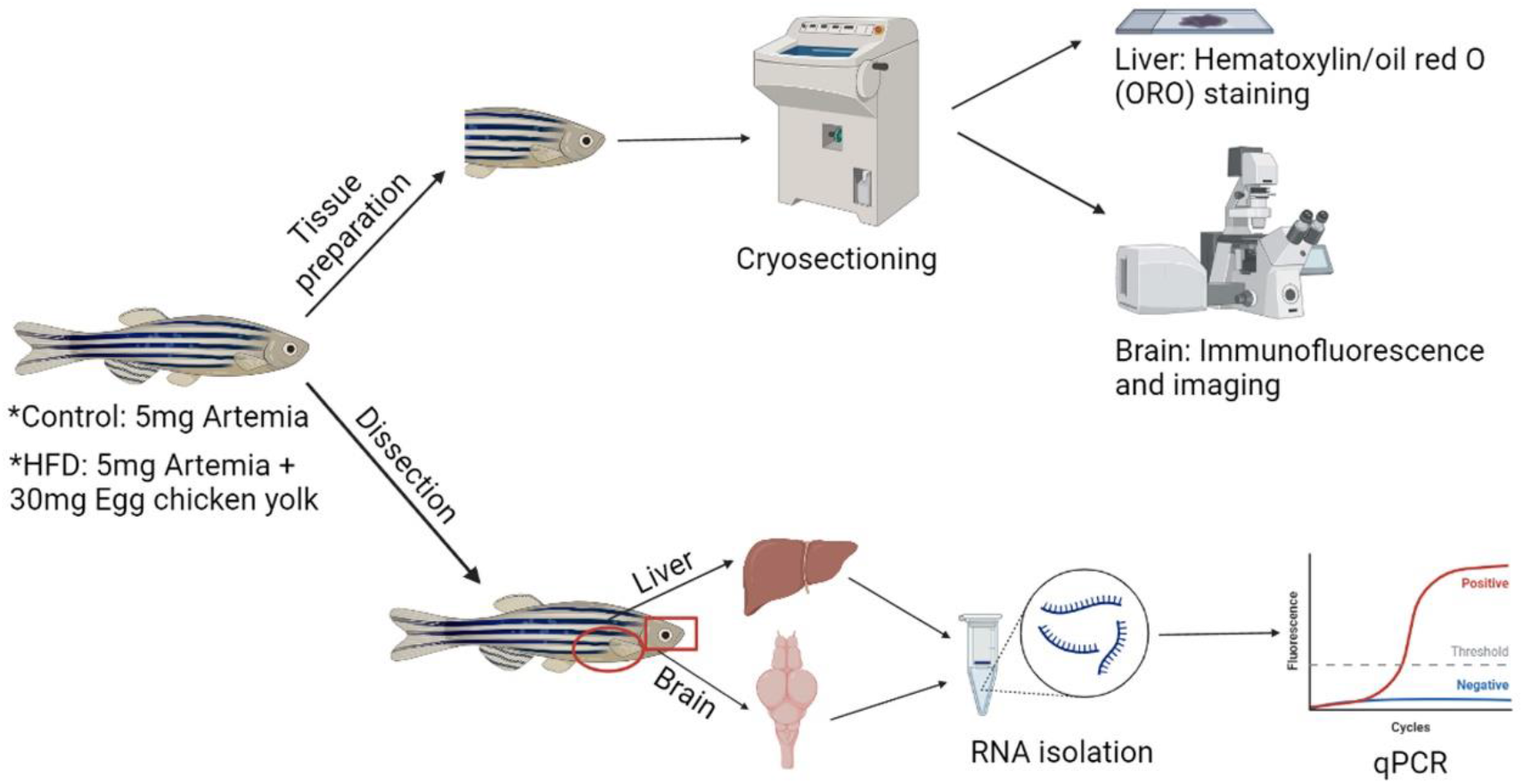
The experimental workflow. The control group of zebrafish were fed with 5 mg of artemia daily. The high-fat diet (HFD) group were fed with 5 mg of artemia and 30 mg of chicken egg yolk daily. Eight fish were used in each group. Half of them were sacrificed for tissue preparation. Cryosections of livers and brains were processed for hematoxylin-oil red O (ORO) staining and immunofluorescence staining, respectively. The other half was used for dissection of the liver and brain, which were further processed for RNA isolation and quantitative PCR (qPCR) analysis of gene expression.

### Tissue preparation and cryosectioning

The zebrafish were sacrificed in ice water, and the severed heads and bodies from four zebrafish were fixed in 4% paraformaldehyde (PFA) in phosphate-buffered saline (PBS) for 2 days at 4°C as described previously (Demirci et al., 2020). The heads were then incubated in 20% sucrose-20% EDTA for 2 days at 4°C and in 30% sucrose-20% EDTA for another 2 days at 4°C. Next, heads were embedded in a solution of 20% sucrose-7.5% gelatin kept in a styrofoam box filled with dry ice, and stored at -80°C until sectioning. Transverse sections of 14 μm were cut from the olfactory bulb, telencephalon, and liver at -25°C using a cryostat (Leica CM1850, Wetzlar, Germany) and stored at -20°C until further use.

### Immunohistochemistry and oil red O Staining

Stock and working solutions of oil red O (ORO) (Sigma Aldrich, MO, USA) were prepared as described previously (Mehlem et al., 2013). Frozen transverse sections of four zebrafish livers were used for ORO staining. Sections stored at -20°C were kept at room temperature (RT) for 30 min. Next, sections were rinsed in 50% isopropanol in distilled water for 30 seconds, stained with freshly prepared oil red O working solution for 5 min, rinsed for 30 seconds in 50% isopropanol, and dipped 5 times in hematoxylin for nuclei staining. Finally, the sections were rinsed in running water for 30 min and mounted in 70% glycerol diluted with PBS. Four different sections with no tissue loss were counted for the quantification of cells. Statistical significance was evaluated using an unpaired t-test.

### Immunofluorescence staining and imaging

Immunofluorescence staining was carried out as previously described (Demirci et al., 2020). The brain sections on slides were dried at RT for 30 min and washed twice in PBSTX (PBS/0.3% Triton X-100) for 10 min at RT. Following a 15-min incubation with 10 mM sodium citrate at 85°C, the sections were washed twice with PBS for 5 min and incubated with the primary antibody in PBSTX overnight at 4°C. The next morning, the slides were rinsed three times with PBSTX for 10 min at RT, incubated with the secondary antibody, and washed three times with PBS for 5 min at RT. Sections were imaged by using an LSM 880 laser scanning confocal microscope (Carl Zeiss AG, Oberkochen, Germany). The primary antibodies are listed as follows: mouse anti-HuC/HuD (1:400, A-21271, Thermo Fisher Scientific, MA, USA), rabbit anti-cleaved-Caspase-3 (1:400, 5A1E, Cell Signaling Technology, MA, USA) mouse anti-PCNA (1:200, M0879, Dako, Agilent, CA, USA), rabbit anti-S100β (1:200, Z0311, Dako, Agilent, CA, USA) and rabbit anti-phospho-β-catenin (Ser675, 1:200, D2F1, Cell Signaling Technology, MA; USA). Secondary antibodies are listed as follows: Rhodamine (TRITC)-AffiniPure donkey anti-rabbit IgG (1:200, 711-025-152, Jackson Immunoresearch Laboratories, PA, USA) and Cy5 AffiniPure donkey anti-mouse IgG (H+L) (1:200, 712-175-150, Jackson Immunoresearch Laboratories, PA, USA). Nuclear staining was carried out by using 4’,6-diamidino-2-phenylindole (DAPI; 4083S, Cell Signaling Technology, MA, USA). Four different sections with no tissue loss were counted for the quantification of cells. Statistical significance was evaluated using an unpaired t-test.

### Western blotting

Four zebrafish from each group (control, 1 month fed, and 2 months fed) were sacrificed according to ethical regulations. Shortly after the sacrification, total brain tissues were dissected and pooled for each group. Collected tissues were flash-frozen in liquid nitrogen and stored at -80°C until cell lysate extraction. For extraction, tissues were homogenized in RIPA Lysis and Extraction Buffer (Thermo Fisher Scientific, MA, USA) and centrifuged at max speed for 30 minutes +4°C. Supernatants were collected and dissolved in 5X SDS loading buffer to be used for western blotting. For western blotting, samples were separated by a 12% acrylamide-bis acrylamide gel and transferred to a polyvinylidene fluoride (PVDF) membrane (GE Healthcare, IL, USA). Blocking was performed with 5% milk powder for 45 min at RT and incubated in the following antibodies at the indicated dilutions. Primary antibodies: rabbit anti-PARP (1:500; 9542T, Cell Signaling Technology, MA, USA) and rabbit anti-β-Actin (1:1000; 8457S, Cell Signaling Technology, MA, USA). Secondary antibodies: anti-rabbit IgG, horseradish peroxidase (HRP)-linked (1:1000; 7074S, Cell Signaling Technology, MA, USA) and goat anti-Rabbit IgG (H+L) Cross-Adsorbed Secondary Antibody, Alexa Fluor™ 546 (1:1000; A-11010, Thermo Fisher Scientific, MA, USA).

### RNA isolation and cDNA synthesis

After 1 month or 2 months of feeding a normal diet or HFD, zebrafish were sacrificed in ice water. Heads were separated from the fish and whole brains were dissected from four zebrafish and pooled. Livers were likewise dissected from four zebrafish and pooled. Total RNAs were isolated from the brain and liver samples using a miRNeasy Mini Kit (Qiagen, Hilden, Germany). cDNAs were synthesized from total RNAs using iScript reverse transcription supermix for RT-qPCR (Bio-Rad Laboratories, Inc., CA; USA) according to the manufacturer’s instructions, using a 1:1 mixture of oligodT and random primers.

### Quantitative PCR (qPCR) and statistical analysis

qPCR was performed in triplicates using the GoTaq qPCR master mix (Promega, Madison, WI, USA) at Applied Biosystems 7500 Fast Real-Time PCR Machine (Foster City, CA, USA) as triplicates. Zebrafish *ribosomal protein L13a* (*rpl13a*) gene was used as a reference for normalization to determine relative gene expression levels. Data were analyzed using the GraphPad Prism 7 software (Graphpad Software Inc., CA, USA). The values are mean ± SEM (Standard Error of Mean) of three samples. Primer sequences for the following zebrafish genes are listed in Table 1: *rpl13a, il6st, il8, mpeg1.1, il2rga, il10, marco, caspase3a, caspase8, PCNA, GFAP, olig2, huD, neuroD1, rbfox3a, axin2* and *lef1.* Statistical significance was evaluated using an unpaired t-test was used for statistical calculations.

**Table 1.**
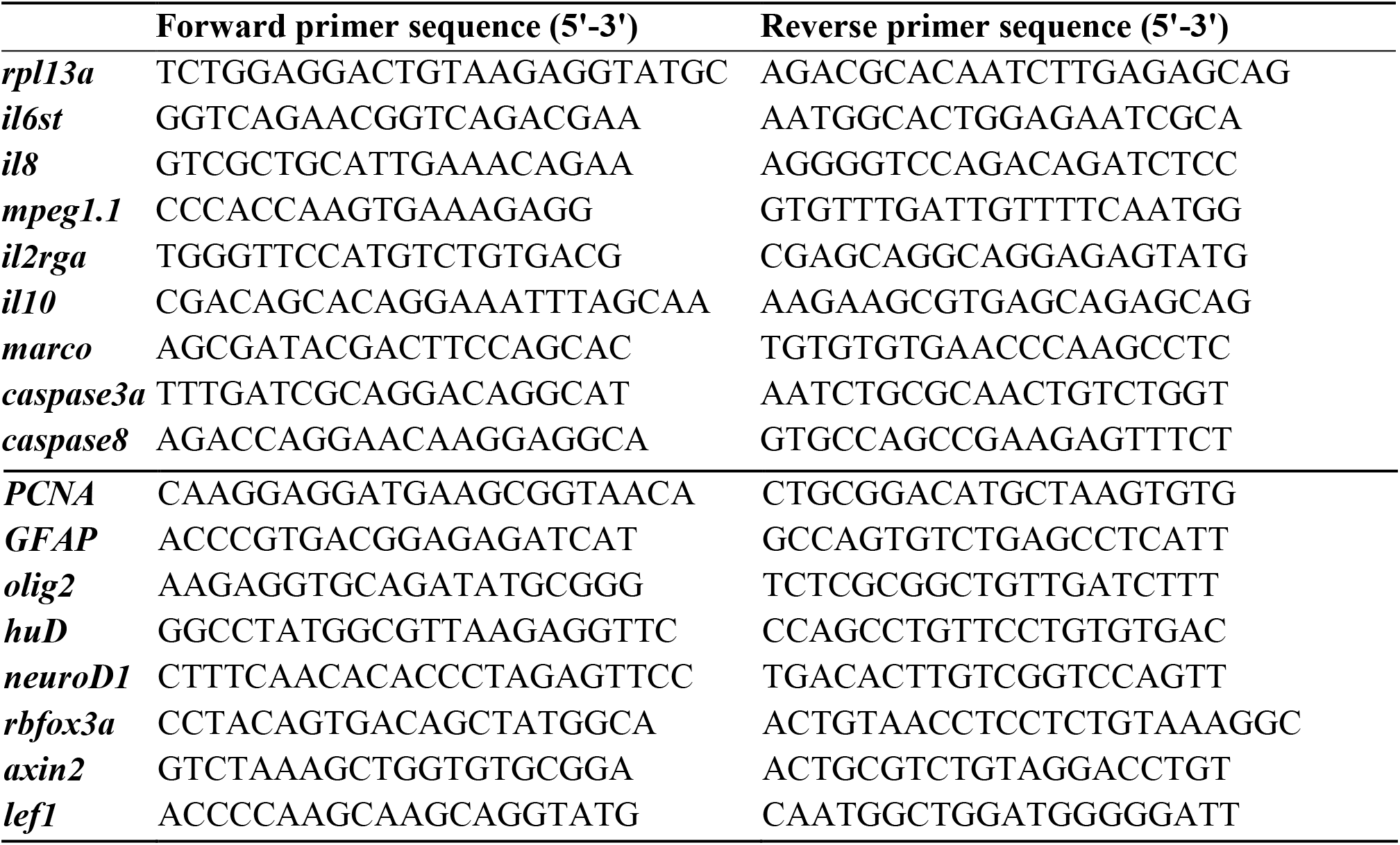
Sequences of forward and reverse primers

### Behavioral assays

Behavioral tests were carried out after the fish (n=7-9 per group) were fed an HFD for 1 or 2 months. The tests were carried out between 10 am and 2 pm using a trapezoidal test tank (LxWxH:27 cm x 10 cm x 17 cm) filled with 3.5 L of system water. After each fish was tested, tank water was replaced with new system water (27 ±1°C). Assay tanks were recorded simultaneously with two different cameras (iPhone 12 Pro Max and iPhone 13). The recordings were then analyzed with Panlab Smart v3.0.

### Novel tank diving assay

The novel tank diving assay, similar to the thigmotaxis test that is used to study anxiety in rodents, is used to measure anxiety in adult zebrafish upon exposure to a new environment. In addition, it is used to measure locomotor activity. The tank was filled with water to a height of 15 cm and divided into 3 horizontal zones of top, middle, and bottom, 5 cm each. The recording was started immediately after the fish were placed in the tank for 1 minute at each of the 0-5, 6-10, 11-15, 16-20, 21-25, and 26-30 minutes intervals. Data were analyzed for the total distance (cm), mean speed (cm/s), distance in the bottom zone (cm), time spent in the bottom zone (s), and the number of entries to the top zone (Audira et al., 2018).

### Mirror-biting assay

The mirror-biting test is a well-established fish paradigm that is widely used to study adult zebrafish’s social and aggressive behavior (Pham et al., 2012; Liu et al., 2020). The test is conducted with a mirror placed on one side of the tank and reflects the reaction of the adult zebrafish to its mirror image. ‘Biting’ is defined as the fish’s “tracing” behavior of its reflection as it swims quickly back and forth, and is measured by the time the fish spends in front of the mirror and touches the mirror (Audira et al., 2018). A mirror was placed at one end of the tank, which was then divided into 2 vertical zones as the mirror-close zone (1 cm from the mirror) and the mirror-far zone (the rest of the tank). The mirror was covered with an opaque plastic sheet to prevent the fish from reacting to their reflections during the acclimatization process. After the fish were placed in the tank, they were allowed to acclimate for 5 minutes. Following the acclimatization, fish behavior was recorded for 15 minutes. Data were analyzed for the total distance (cm), mean speed (cm/s), distance in the mirror-close zone (cm), time spent in the mirror-close zone (s), and the number of entries to the mirror-close zone. calculating the Distance in the mirror-close zone (cm), total distance (cm), number of entries in the mirror-close zone, time spent in the mirror-close zone (s), and the mean speed (cm/s) (Audira et al., 2018).

## Results

### High-fat diet induces weight gain and lipid accumulation in the liver

To validate the influence of HFD feeding we dissected the livers from 1 month and 2 months fed zebrafish. The fat build-up in the liver was considerably higher in fish fed an HFD for 2 months (long-term) than for 1 month (short-term) (Figure 2A). Consistent with the fat accumulation, there was a significant and gradual increase in the average body-mass index (BMI) of the fish (Figure 2B). The number and size of the lipid droplets harbored in the liver increased in fish fed HFD, as detected by hematoxylin-oil red O (ORO) staining (Figure 2C–E). These results confirm that HFD leads to an increase in body weight and the build-up of excess fat in the liver.

**Figure 2.**
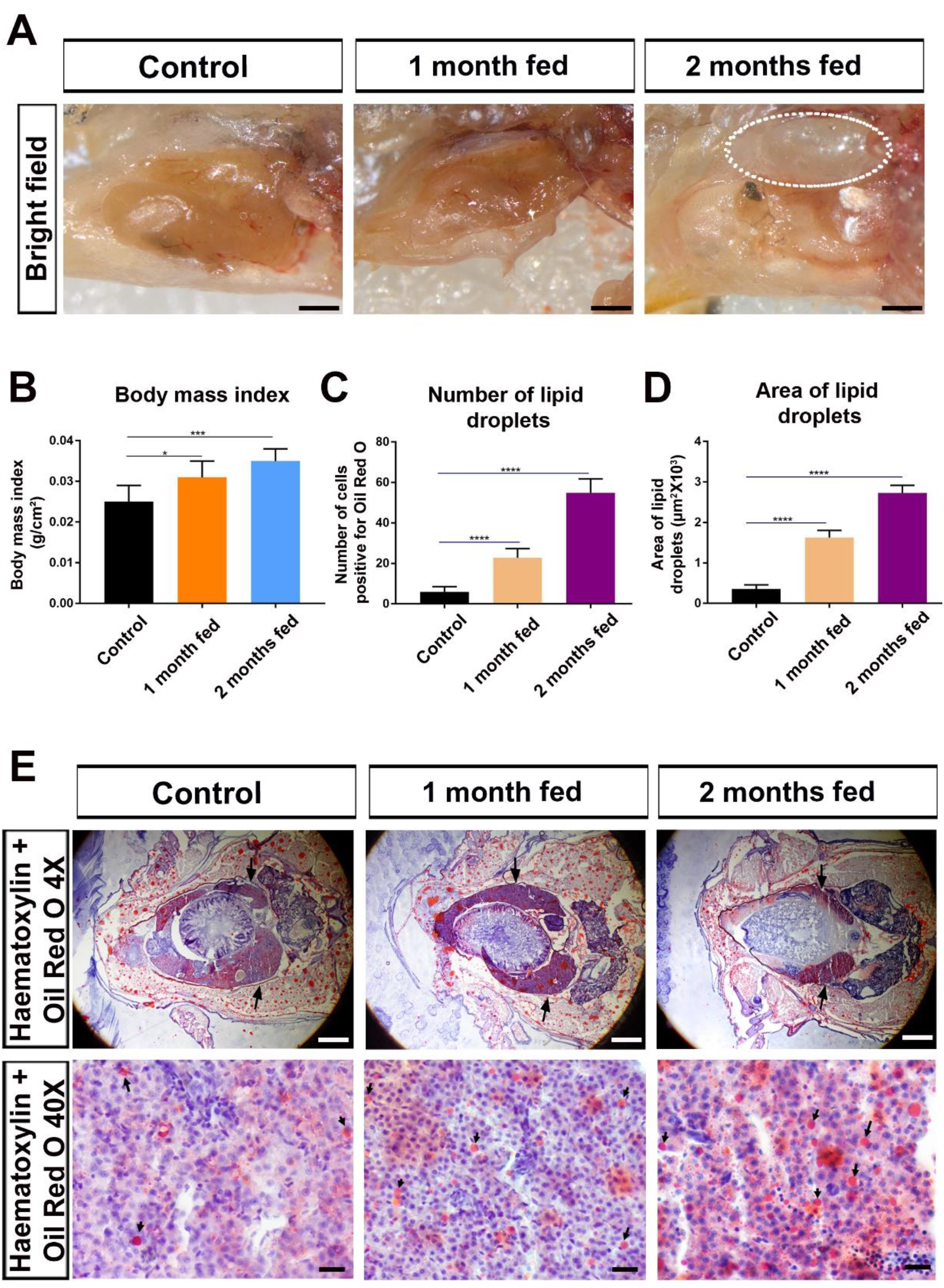
High-fat diet induces weight gain and lipid accumulation in the liver. **(A)** Bright-field images display lipid build-up in the livers of zebrafish fed an HFD most prominently after 2 months of feeding. Dotted lines indicate large fat deposits in the liver. **(B)** The body-mass index (BMI) graph of zebrafish increased significantly after 1 and 2 months of HFD as compared to the control group. **(C)** The number of cells positive for Oil Red O in the liver of control, 1-month fed, and 2-month fed zebrafish. Statistical significance was evaluated using an unpaired t-test. ****p<0.0001. Error bars represent ± standard error of the mean (SEM, n=3). **(D)** The area of lipid droplets in the liver of control, 1-month-fed, and 2-month-fed zebrafish. Statistical significance was evaluated using an unpaired t-test. ****p<0.0001. Error bars represent ± standard error of the mean (SEM, n=3). **(E)** Hematoxylin (blue-purple) and oil red O (ORO, red) staining of liver sections obtained from 1-month fed and 2 months-fed zebrafish show an increase in the number and size of the lipid droplets at 4X and 40X magnification. Arrows indicate the liver and the lipid droplets in 4X and 40X, respectively. Please note that the overall number and size of the lipid droplets increase in the HFD groups. Scale bars 5 mm in A and upper row of D, 20 μM in the lower row of E.

### HFD causes activation of inflammation, apoptosis, proliferation, and neurogenesis

HFD has been shown to induce the clustering of macrophages that release pro-inflammatory cytokines and chemokines in the zebrafish larval liver (de Oliveira et al., 2019; Shwartz et al., 2019). Activation of the immune response also plays a critical role in the promotion of neural stem cell proliferation and neuronal regeneration (Kyritsis et al., 2012; Kizil et al., 2015). To test whether inflammation response is activated in response to HFD, we measured the expression levels of several immune response marker genes in the liver and brain tissues. We observed that the pro-inflammatory cytokine receptor *interleukin 6 cytokine family signal transducer* (*il6st/glycoprotein 130*), the cytokine *il8,* and the macrophage marker *mpeg* significantly increased in the livers of the fish fed an HFD for short-or long-term (Figure 3A). Activation of the immune response was accompanied by an increase in the apoptotic markers *caspase8* (*casp8*) and *caspase3a* (*casp3a*) as well as in the proliferation marker *proliferating cell nuclear antigen (PCNA)* (Figure 3B). We found a similar elevation in the levels of the pro-inflammatory markers *il2rga* and *marco* and the macrophage marker *mpeg* with a concomitant decrease in the anti-inflammatory marker *il10* in the brain tissues after long-term HFD (Figure 3C). Apoptosis-related genes *casp8* and *casp3a* and *PCNA* were significantly upregulated after short- and long-term feeding (Figure 3D). There was a concomitant increase in the expression of the glial marker *glial fibrillary acidic protein* (*GFAP*), which is known to be strongly upregulated in response to traumatic brain injury (Figure 3E, (Kroehne et al., 2011)). Interestingly, there was a decrease in the oligodendrocyte marker *oligodendrocyte transcription factor* (*olig2*) and the neuronal markers *neuronal differentiation 1* (*neuroD1*), *ELAV-like neuron-specific RNA binding protein 4* (*HuD*), and *Rbfox3a* (*neun*) in response to short-term HFD and a remarkable recovery after long-term HFD, proposing that the dying oligodendrocytes and neurons were replaced with the newborn ones (Figure 3E). Since these genes are expressed in mature neurons in the adult zebrafish brain (Kroehne et al., 2011; Dohaku et al., 2019; Erbaba et al., 2020; Shimizu et al., 2021), our data suggest that HFD results in neuronal cell death, glial cell activation and induction of a regenerative response in the brain. Together, these data indicate that HFD triggers inflammation, apoptosis, and proliferation not only in the liver but also in the brain and reactive neurogenesis in the brain.

**Figure 3.**
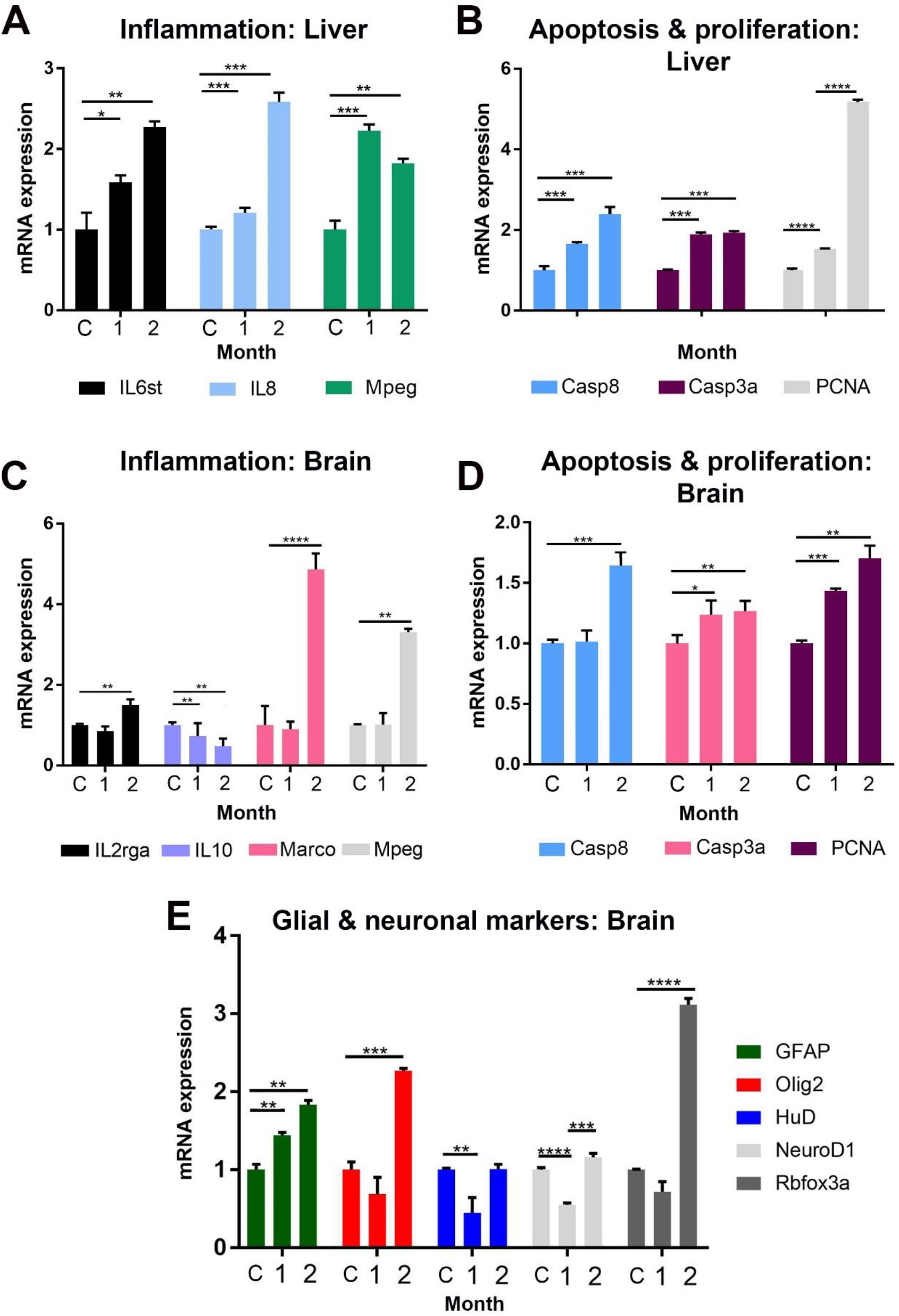
HFD causes activation of inflammation, apoptosis, proliferation, and neurogenesis in the liver and the brain. Expression levels of **(A)** inflammation-related genes in the liver **(B)** apoptosis- and proliferation-related genes in the liver, **(C)** inflammation-related genes in the brain, **(D)** apoptosis- and proliferation-related genes in the brain and **(E)** glial and neuronal markers in the brain determined by qPCR in 1 month fed and 2 months fed fish relative to control fish fed a normal diet. Statistical significance was evaluated using an unpaired t-test. *p<0.05, **p<0.01, ***p<0.001 and ****p<0.0001. Error bars represent ± standard error of the mean (SEM, n=3). Three independent experiments were conducted. C: control.

### Long-term exposure to HFD stimulates apoptotic cell death, glial cell activation, and proliferation in the adult brain

Since expression of apoptosis-related genes in the brain was downregulated after 1 month of HFD feeding and upregulated after 2 months of feeding, next, we aimed to examine the detailed response of different brain regions to HFD feeding concerning the activation of apoptosis. Immunofluorescence staining for cleaved-Caspase 3 (C-Caspase3) showed that apoptosis increased in the olfactory bulb after short-term feeding and was further boosted after prolonged feeding (Figures 4A compare middle and right panels to left panel, 4C). A parallel upregulation of C-Caspase3 was detectable in the telencephalon after 2 months of feeding (Figure 4B–C). Cleaved PARP, another marker of apoptosis, also increased in the total brain extracts of zebrafish fed a long-term HFD (Figure 4D).

**Figure 4.**
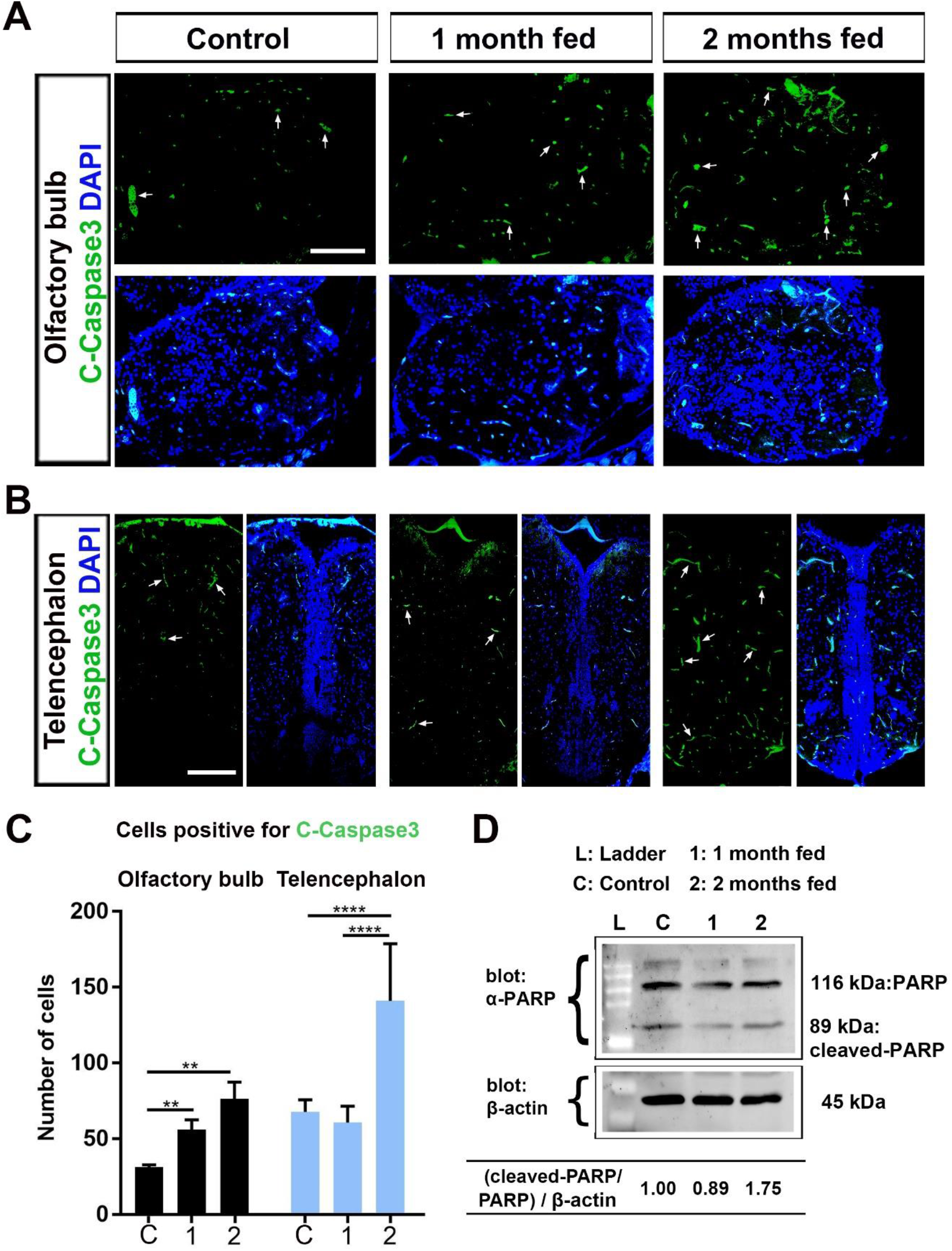
Long-term exposure to HFD stimulates apoptotic cell death in the adult brain. Anti-cleaved-Caspase-3 (green) staining of brain sections obtained from the (**A**) olfactory bulb and **(B)** telencephalon regions of control, 1 month fed, and 2 months fed zebrafish. Arrows indicate several apoptotic cells. Sections are counterstained for DAPI. Scale bars 200μm. **(C)** The number of cells positive for C-Caspase3 in the brain sections obtained from the olfactory bulb and telencephalon regions of control (C), 1 month fed and 2 months fed zebrafish. Statistical significance was evaluated using an unpaired t-test. **p<0.01 and ****p<0.0001. Error bars represent ± standard error of the mean (SEM, n=3). **(D)** Western blot of whole brain tissue lysates collected from control, 1 month fed and 2 months fed to fish. Cleaved-PARP/PARP to β-actin ratios were calculated.

Activation of RGCs upon injury is a hallmark of regeneration in the adult zebrafish brain (Jurisch-Yaksi et al., 2020). RGCs are characterized by a few markers including S100β and express the cell proliferation marker PCNA. To unravel the regenerative response of RGCs to HFD feeding, we performed immunofluorescence staining of the proliferating RGCs for S100β and PCNA. We detected a gradual increase in glial cell proliferation, marked by S100β and PCNA double-positive cells, which became prominent in the olfactory bulb and telencephalon after HFD feeding (Figure 5A–B). The number of double-positive cells in both CNS tissues increased concurrently, marking a substantial part of the glial cells located at the periventricular zones (Figures 5A–B compare green and red cells in boxes, Figure 5C). Strikingly, proliferating cells marked by PCNA were not restricted to the ventricular zones in the telencephalon but were also detected deeper in the brain parenchyma (Figure 5B). These results collectively suggest that brain tissue is highly responsive to HFD feeding concerning activation of apoptosis, glial cell response, and proliferation.

**Figure 5.**
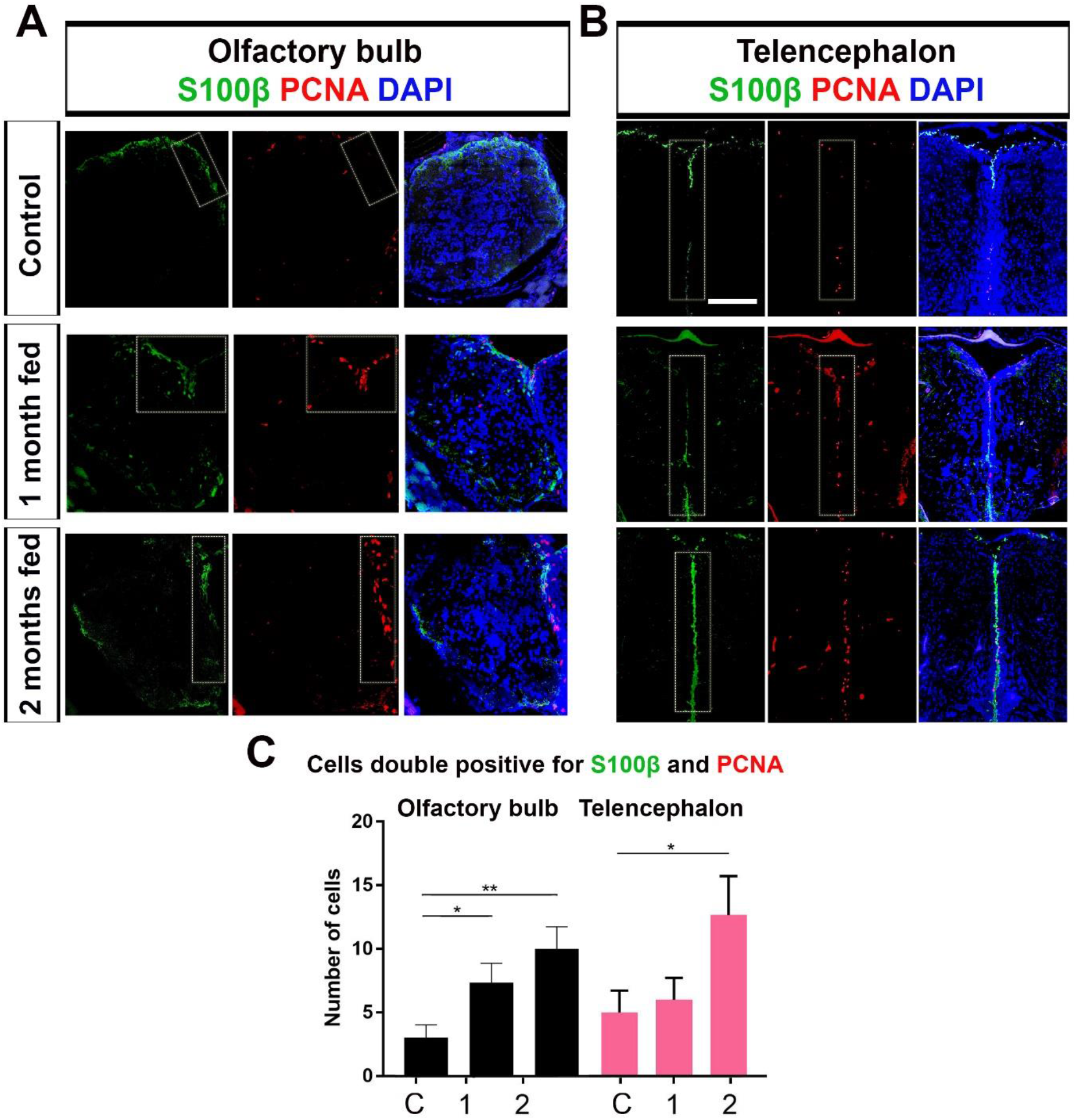
Long-term exposure to HFD stimulates glial cell activation and proliferation. Anti-s100β (green) and anti-PCNA (red) staining of brain sections obtained from the **(A)** olfactory bulb and **(B)** telencephalon of control, 1 month fed, and 2 months fed zebrafish. Dotted rectangles indicate periventricular zones where radial glial cells proliferate. Dotted ellipses indicate brain parenchyma. Sections are counterstained for DAPI. Scale bars 200μm. **(C)** The number of cells double positive for S100β and PCNA in the brain sections obtained from the olfactory bulb and telencephalon regions of control (C), 1 month fed and 2 months fed zebrafish. Statistical significance was evaluated using an unpaired t-test. *p<0.05 and **p<0.01. Error bars represent ± standard error of the mean (SEM, n=3).

### HFD intake results in initial loss and subsequent recovery of neurons

An increase in apoptosis and concomitant reduction of mRNA levels of the neuronal markers *neuroD1* and *HuD* in the brain suggests that HFD could promote neuronal cell death. To test this hypothesis, we examined the expression of HuC/D by antibody staining on the sections of the olfactory bulb and the telencephalon obtained from 1 month and 2 months fed zebrafish. There was a significant reduction in neuron densities in the olfactory bulb of the fish fed a short-term HFD as compared to the control (Figure 6A middle column, Figure 6C). However, long-term HFD resulted in a considerable recovery in the number of HuC/D-positive neurons (Figure 6A right column, Figure 6C). Strikingly, following a similar decrease at 1 month of HFD, the neuron densities were substantially restored in the telencephalon at 2 months of HFD (Figure 6B compare middle columns to right columns, Figure 6C). Thus, short-term HFD induces neuron loss, which is compensated by the new neurons emerging in the course of prolonged exposure to HFD.

**Figure 6.**
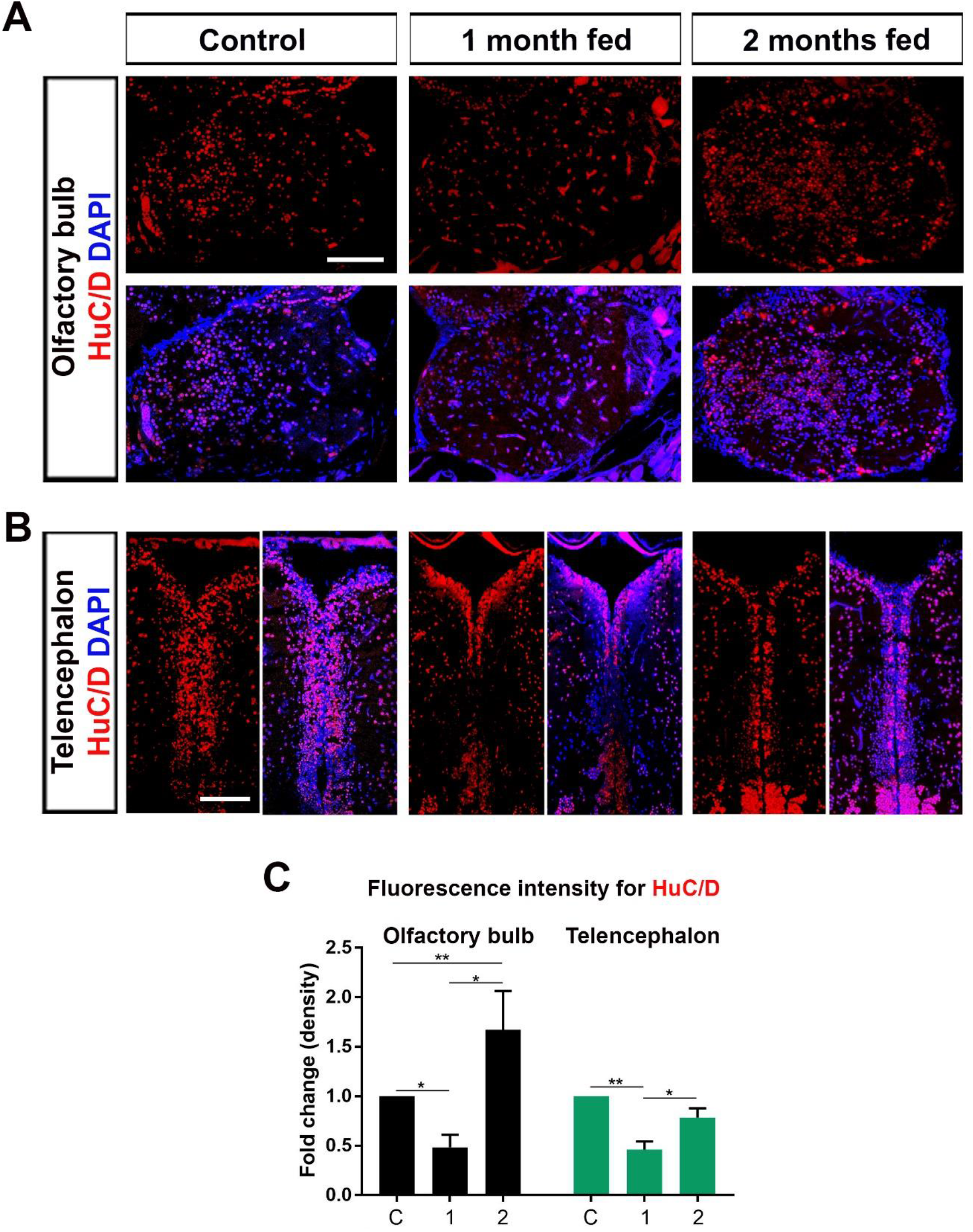
HFD intake results in loss and subsequent recovery of neurons in the brain. Anti-HuC/D (green) staining of brain sections obtained from the **(A)** olfactory bulb **(B)** telencephalon of control, 1 month fed, and 2 months fed zebrafish. Sections are counterstained for DAPI. Scale bars 200μm. (C) The fluorescence intensity of cells positive for HuC/D in the brain sections obtained from the olfactory bulb and telencephalon regions of control **(C)**, 1 month fed and 2 months fed zebrafish. Statistical significance was evaluated using an unpaired t-test. *p<0.05 and **p<0.01. Error bars represent ± standard error of the mean (SEM, n=3).

### HFD feeding activates Wnt/β-catenin signaling in the brain

Wnt/β-catenin signaling pathway becomes activated in the regenerating zebrafish telencephalon at the early wound healing stage, supporting the tissue-wide regeneration-promoting role of the pathway (Demirci et al., 2020). Immunofluorescence staining of the olfactory bulb and the telencephalon revealed elevation of phospho-β-catenin (phosphorylated at Ser675, increasing nuclear localization and transcriptional activation of β-catenin) in the brains of the fish fed HFD for short-term and a further increase in response to prolonged HFD feeding (Figure 7A–B). The increase was reflected in the number of cells positive for β-catenin (Figure 7C). Finally, qPCR results confirmed canonical Wnt signaling activation as evidenced by significant upregulation of the Wnt/β-catenin target genes *axin2* and *lef1,* as well as β-catenin in the total brain tissues of fish, fed an HFD for 2 months (Figure 7D). Thus, we conclude that Wnt/β-catenin signaling is activated in the brain in response to HFD.

**Figure 7.**
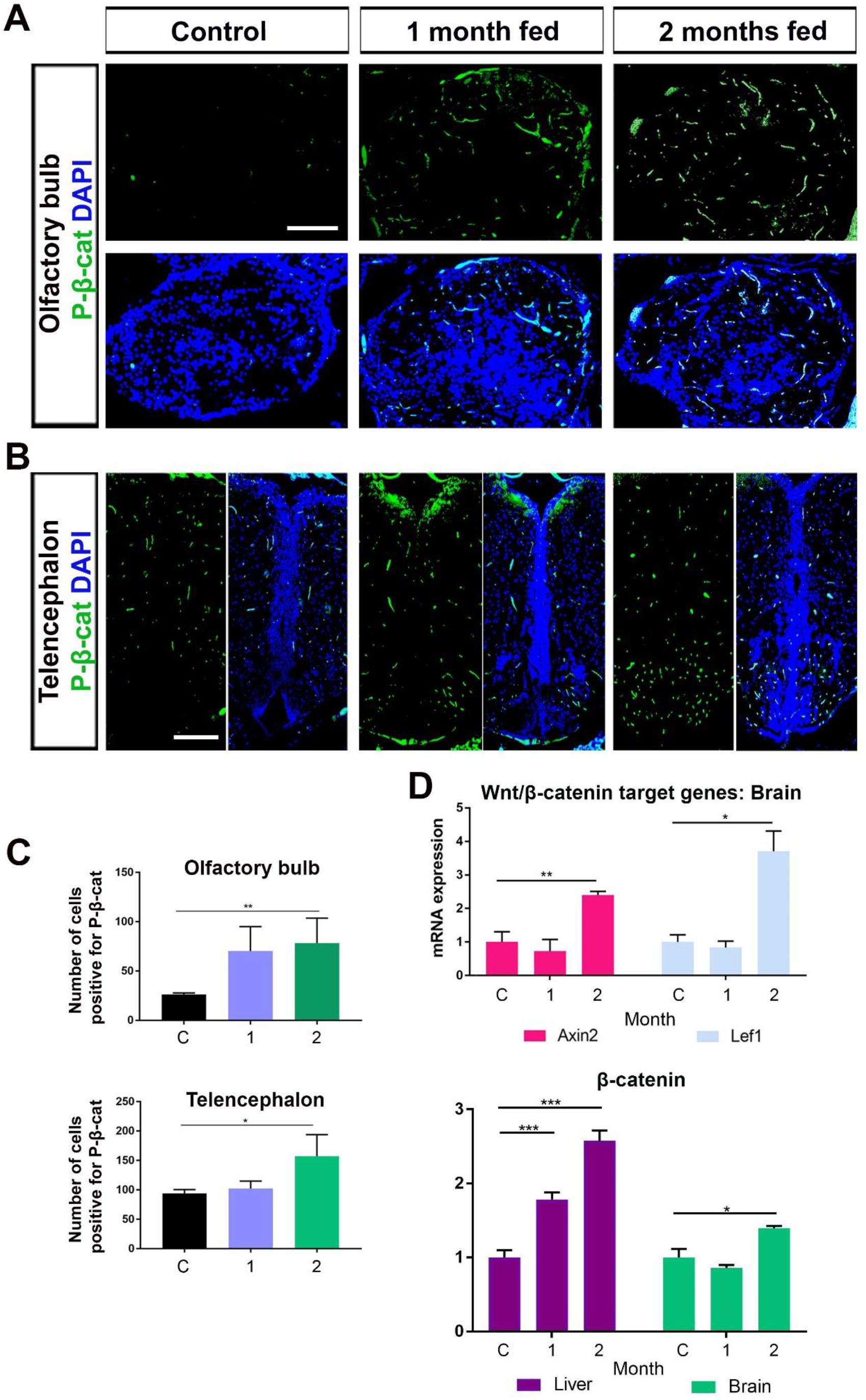
HFD feeding activates Wnt/β-catenin signaling in the brain. Anti-phospho-β-catenin (green) staining of brain sections obtained from the **(A)** olfactory bulb and **(B)** telencephalon of control, 1-month fed, and 2-month fed to zebrafish. Sections are counterstained for DAPI. Scale bar=100μm. **(C)** The number of cells positive for phospho-β-catenin in the brain sections obtained from the olfactory bulb and telencephalon regions of control (C), 1 month fed and 2 months fed zebrafish. Statistical significance was evaluated using an unpaired t-test. *p<0.05 and **p<0.01. Error bars represent ± standard error of the mean (SEM, n=3). **(D)** Relative expression levels of canonical Wnt target genes Axin2 and Lef1 in the brain were determined by qPCR in 1-month-fed and 2-months-fed fish relative to control (C) fish fed a normal diet. Statistical significance was evaluated using an unpaired t-test. *p<0.05, **p<0.01 and ***p<0.001. Error bars represent ± standard error of the mean (SEM, n=3). Three independent experiments were conducted.

### HFD feeding causes an increase in anxiety, aggressiveness, and locomotor activity in zebrafish

HFD feeding is known to exert neurobehavioral effects including anxiety/depressive-like behaviors and aggressiveness (Baker and Reichelt, 2016; de Noronha et al., 2017; Picolo et al., 2021). Similar to most other species, exposure to novelty gives rise to robust anxiety responses in zebrafish. Thus, zebrafish exhibit bottom-dwelling behavior when placed in a new environment and begins to explore the area only after a few minutes of adaptation (Egan et al., 2009; Parker et al., 2014). To explore whether HFD feeding influences the anxiety level of the zebrafish, we used the novel tank diving assay, a behavioral test based on measuring the instinctive behavior of zebrafish when placed in an unfamiliar environment as well as the locomotor activity. First, the total distance traveled in the first 15 minutes and mean speed in this period significantly increased in fish fed with an HFD for 1 or 2 months compared to the control (Figure 8A–B). These results show that HFD feeding results in an early increase in locomotor activity. Moreover, fish fed an HFD for short- or long-term traveled longer distances and spent significantly longer time at the bottom zone of the tank, and displayed longer latency to enter the top zone in almost all periods (Figure 8C–D). In several periods including the first 5 minutes, the number of entries to the top zone of the tank also decreased in HFD groups compared to the control (Figure 8E).

**Figure 8.**
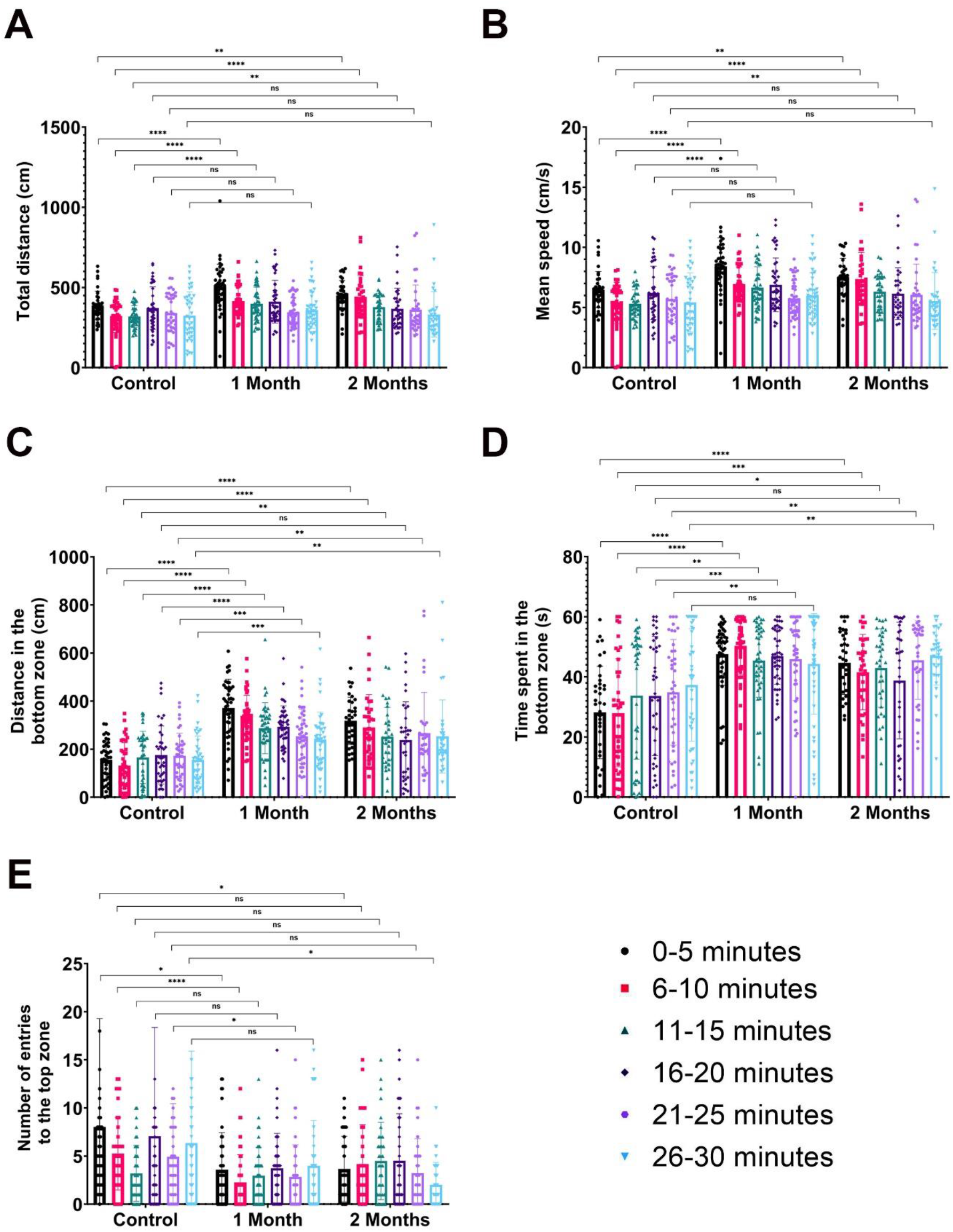
HFD feeding causes an increase in anxiety in zebrafish. (A) Total distance (cm) (B) mean speed (cm/s), (C) distance in the bottom zone (cm), (D) time spent in the bottom zone (s) and (E) number of entries to the top zone measured for fish fed a control diet or HFD for 1 month or 2 months. Statistical significance was evaluated using an unpaired t-test. *p<0.05, **p<0.01, ***p<0.001, ****p<0.0001, and ns nonsignificant. Error bars represent ± standard deviation of the mean (SD, n=7-9).

Next, to test whether HFD feeding induces any aggressive behavior, we exploited the mirror-biting assay, which is used to assess the level of aggressiveness in the social behavior of adult zebrafish (Cachat et al., 2013; Liu et al., 2020). We observed a significant increase in the total distance traveled and mean speed of the fish fed an HFD for 1 or 2 months compared to the control, similar to the novel tank diving assay (Figure 9A–B). Short- or long-term HFD feeding led to an increase in the distance traveled, but not time spent, in the mirror-close zone (Figure 9C–D). Besides, short-term HFD-fed fish displayed a tendency to enter more frequently in the mirror-close zone (Figure 9E). These data together indicate that HFD feeding likely triggers anxiety-like behavior and aggressiveness in zebrafish, evidenced by delayed adaptation to the new environment and increased tendency to attack their reflection in the mirror, respectively.

**Figure 9.**
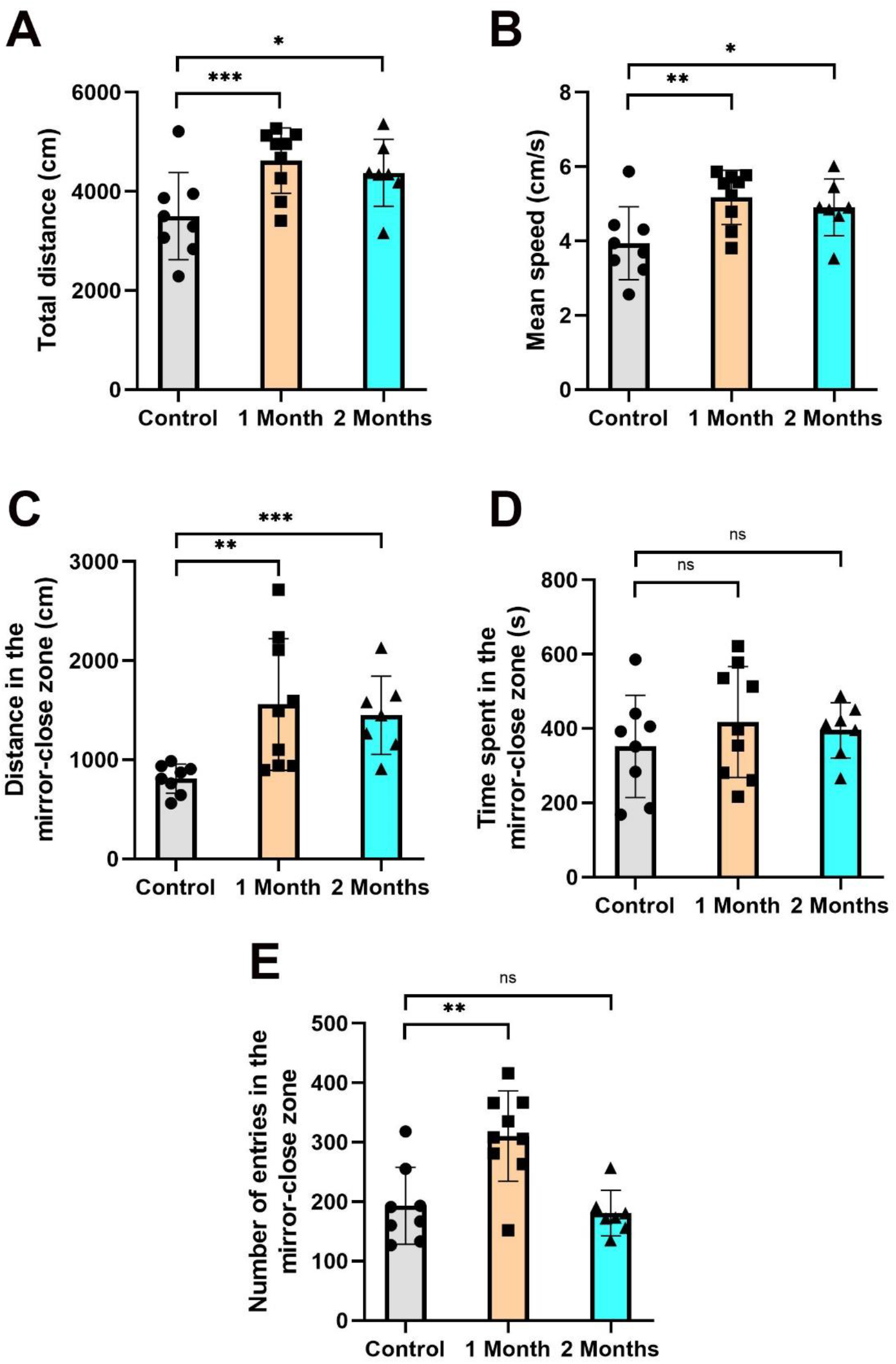
HFD feeding causes an increase in aggressiveness in zebrafish. (A) Total distance (cm), (B) mean speed (cm/s), (C) distance in the mirror-close zone (cm), (D) time spent mirror-close zone, and (E) number of entries in the mirror-close zone measured for fish fed a control diet or HFD for 1 month or 2 months. Statistical significance was evaluated using an unpaired t-test. *p<0.05, **p<0.01, ***p<0.001, and ns nonsignificant. Error bars represent ± standard deviation of the mean (SD, n=7-9).

## Discussion

NAFLD is characterized by the build-up of excess fat in the liver cells that is not caused by alcohol and is mostly linked to HFD feeding-induced obesity. NAFLD can have a systemic effect on other organs over time. In this study, we used an adult zebrafish model of HFD feeding to unravel how HFD affects the brain. Our data show that HFD feeding i) is detectable as weight gain and abnormal lipid deposition in the liver, ii) is accompanied by an increase in the expression levels of the genes associated with inflammation, apoptosis, and proliferation in the liver and brain tissues, iii) stimulates glial cell activation and neuronal turnover in the forebrain (olfactory bulb and telencephalon), iv) activates canonical Wnt signaling pathway in these brain regions and v) increases anxiety, aggressiveness and locomotor activity (Figure 10).

**Figure 10.**
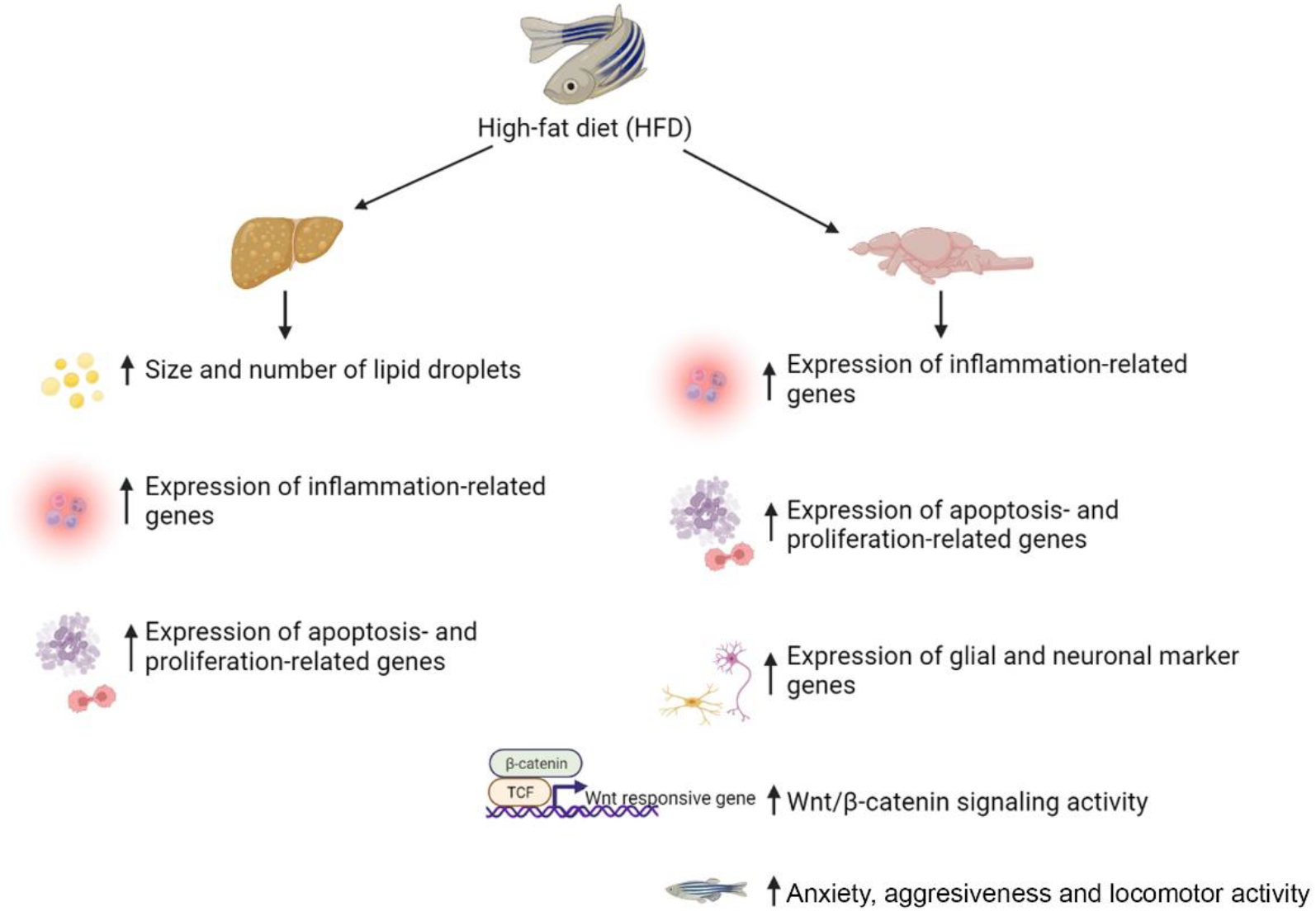
Summary of the influence of HFD feeding in the liver and the brain. (Left) HFD increases the size and number of lipid droplets, the elevation of inflammation-, apoptosis- and proliferation-related genes’ expression. (Right) HFD feeding leads to changes in the expression of genes related to inflammation, apoptosis, proliferation, neurogenesis, and Wnt/β-catenin signaling pathway in the brain as well as neurobehavioral effects.

Zebrafish constitute a useful model to study mammalian diet-related pathological conditions as HFD elicits similar biological responses in mammals and zebrafish (Vargas and Vásquez, 2017; Carnovali et al., 2018; Meguro et al., 2019; Ka et al., 2020; Picolo et al., 2021). Our results showed that, in parallel to the considerable increase in body weight, fish fed a long-term HFD exhibited fat accumulation in the liver, similar to obesity phenotypes (Oka et al., 2010; Leibold and Hammerschmidt, 2015; Landgraf et al., 2017). In addition to its adverse effects on the liver (Lian et al., 2020), dietary changes including HFD have been associated with brain disorders including dementia, Alzheimer’s disease, and Parkinson’s disease (Morris et al., 2010; Pistell et al., 2010; Petrov et al., 2015; Nuzzo et al., 2018). Thus, our results further support the relationship between HFD-induced obesity and the brain, by showing for the first time that HFD feeding induces a regeneration response accompanied by activation of the canonical Wnt signaling pathway in the brain of the highly regenerative zebrafish.

HFD-induced obesity and NAFLD have been revealed to cause hepatic inflammation (Fabbrini et al., 2010; Heinrich et al., 2015; Chen et al., 2018; Luo and Lin, 2021). Our data showed gradually increasing upregulation of the pro-inflammatory cytokines and their receptors *il6st* and *il8* in the liver and a relatively mild upregulation of *il2rga* and *il6st* (data not shown but similar to *il2rga*) in the brain, in response to prolonged HFD. IL-6 has been identified as a regulatory factor that plays a role in blood coagulation and platelet activation in zebrafish and mammalian obesity (Oka et al., 2010). IL-6 levels are significantly increased in the hippocampus of rats fed an HFD (Dutheil et al., 2016). IL-6 receptor IL6st, a member of the Janus kinase 1/Signal transducer and activator of transcription 3 (Jak1/Stat3) pathway is activated in an injury-dependent manner during regeneration of the heart, optic tectum, retina, and telencephalon (Fang et al., 2013; Zhao et al., 2014; Demirci et al., 2020; Shimizu and Kawasaki, 2021). Another pro-inflammatory marker IL-2R has been found to increase in neuropsychiatric disorders including schizophrenia, depression, and post-traumatic stress disorder (Pape et al., 2019). IL-8 and Marco are also essential for the modulation of the innate immune response during neuronal injury and neuroinflammation (Kossmann et al., 1997; Kushi et al., 2003; Milne et al., 2005; Orr et al., 2017; Du et al., 2018). The macrophage-expressed gene *mpeg*, which is upregulated during bacterial infection, is likewise upregulated during the regenerative response in the liver and the central nervous system (Benard et al., 2015; Tsarouchas et al., 2018; Shwartz et al., 2019; Demirci et al., 2020). On the other hand, high BMI levels have been associated with reduced levels of the anti-inflammatory cytokine IL-10 in the human frontal cortex (Lauridsen et al., 2017). Thus, differential regulation of these early immune response elements upon HFD feeding suggests that they coordinate activation of an injury-induced inflammatory response in both the liver and the brain. Several studies in rodents assert that consumption of an HFD is sufficient to induce an inflammatory response in the brain and increase the permeability of the blood-brain barrier (BBB), disrupting its integrity, especially in the hippocampus, and leading to impaired cognition (De Souza et al., 2005; Kanoski et al., 2010; Pistell et al., 2010; Freeman et al., 2014; Nakandakari et al., 2019; Leigh and Morris, 2020; de Paula et al., 2021). The enhanced permeability of the BBB allows the passage of reactive oxygen species and pro-inflammatory cytokines from the blood to the brain, leading to neuroinflammation (Freeman et al., 2014; Więckowska-Gacek et al., 2021). Therefore, it is likely that the inflammatory response caused by HFD in the liver, results in neuroinflammation triggered by non-traumatic brain damage.

Resolution of inflammation is essential for the restoration of tissue integrity and function and includes apoptosis as a key component to stop inflammation and initiate the healing process (Ortega-Gómez et al., 2013; Wu and Chen, 2014). We observed transcriptional activation of apoptosis-related genes in the liver after short-term HFD and in the brain after long-term HFD. Increased levels of apoptosis in the liver have been observed in liver injury models generated via feeding HFD (Wang et al., 2008; Jia et al., 2020). High fat consumption is also associated with the induction of apoptosis-related genes such as Bcl2 and Bax in the mammalian hippocampus (Nakandakari et al., 2019; de Paula et al., 2021). We detected cleaved-Caspase-3 positive apoptotic cells in the olfactory bulb and telencephalon of the forebrain. The distribution of apoptotic cells in the glomerular layer of the olfactory bulb is consistent where mitral cells are abundant (Fuller et al., 2006). Mitral cells have been reported to be sensitive to metabolic changes such as HFD-induced obesity (Prud’homme et al., 2009; Fadool et al., 2011). Moreover, cleaved-Caspase-3-positive cells in the telencephalon were concentrated in the parenchyma, suggesting that they correspond to not only the neurons but also the oligodendrocytes, endothelial cells or macrophage/microglia (Kroehne et al., 2011). Our results are in line with a previous study showing enhanced expression of *casp9* in the telencephalon of zebrafish fed an HFD (Meguro et al., 2019). Strikingly, both transcriptome analyses of brain tissues and histological examination of brain sections have revealed activation of apoptosis signaling pathways at different stages of regeneration (12 hours post-lesion (hpl), 20 hpl, and 3 days-post lesions) after telencephalic and optic tectum injuries (Kroehne et al., 2011; Demirci et al., 2020; Shimizu et al., 2021; Demirci et al., 2022). Thus, the concomitant increase of inflammatory signals and apoptotic gene expression in our HFD model could point out that the brain gives similar molecular responses to long-term HFD feeding and traumatic brain injury to initiate a healing process. It is also noteworthy that the inflammation response of the whole brain and the apoptotic response of the three different brain tissues is indistinct after short-term HFD feeding but becomes pronounced after long-term HFD feeding, suggesting that neuroinflammation and neural apoptosis are activated in response to prolonged feeding (Van Dyken and Lacoste, 2018).

A key reaction of the zebrafish brain to inflammation is the activation of RGCs, which upregulate proliferation and result in increased neurogenesis (Kroehne et al., 2011; Kyritsis et al., 2012; Demirci et al., 2020; Kanagaraj et al., 2022). Reactive gliosis marked by elevated GFAP expression in astrocytes has also been revealed to accompany brain inflammation as a response to HFD consumption in mouse and rat models (Pistell et al., 2010; Bondan et al., 2019; Micioni Di Bonaventura et al., 2020). These observations are in coherence with our findings where expression of the RGC markers GFAP and S100β and the proliferation marker PCNA were enhanced in different regions of the brain after short- or long-term HFD feeding. Proliferating RGCs were mostly detected at the periventricular zones in the telencephalon. Most strikingly, we found a significant downregulation of the neuronal markers NeuroD1 and HuC/D after short-term HFD consumption and a recovery in their expression after prolonged feeding. This proposes that HFD feeding induces neuronal cell death in the early stage and the newborn neurons replenish the lost neurons over time. Collectively, similar to a traumatic brain injury, HFD stimulates glial cell proliferation and neurogenesis in the adult zebrafish brain.

The Wnt signaling pathway is involved in brain development and function with key roles in the regulation of neurogenesis, synaptogenesis, synaptic plasticity, and neuronal plasticity/repair (Oliva et al., 2013; Rosso and Inestrosa, 2013; McLeod and Salinas, 2018; Arredondo et al., 2020). Moreover, Wnt signaling is deciphered to play a key role in the activation of the innate immune system and neuroinflammation in neurodegenerative diseases and mental disorders (Zolezzi and Inestrosa, 2017; Bem et al., 2019; Palomer et al., 2019; Zhang et al., 2019; Karabicici et al., 2021). Wnt/β-catenin signaling is activated in the early stages of brain regeneration, with a regeneration-promoting role (Salehi et al., 2018; Zhang et al., 2018; Chang et al., 2020; Demirci et al., 2020; Menet et al., 2020). Our results display a prominent activation of Wnt/β-catenin signaling in the total brain tissue as well as in the olfactory bulb and the telencephalon in the brain, after prolonged exposure to HFD. To our knowledge, this is the first evidence of active canonical Wnt signaling in the brain in response to HFD feeding. Thus, Wnt/β-catenin signaling appears to be a primary pathway with a key role in the regulation of the regenerative response of the brain to different types of damage including traumatic brain injury and HFD feeding-induced brain damage.

In humans, obesity and overweight are associated with aggressive behaviors and anxiety/depression (Cerniglia et al., 2018; Lindberg et al., 2020). HFD feeding of mice, rats, or zebrafish results in a similar increase in aggression and anxiety (Buchenauer et al., 2009; Baker and Reichelt, 2016; de Noronha et al., 2017; Picolo et al., 2021). We observed a similar increase in the anxiety and aggressiveness of zebrafish fed an HFD for the short- or long-term. Interestingly, there was a parallel increase in the locomotor activity of fish fed an HFD, shown by both the novel tank diving and mirror-biting assays. Picolo et al. have recently reported that HFD feeding for the short term did not affect the locomotor activity in adult zebrafish (Picolo et al., 2021). However, those experiments were terminated on the 16^th^ day of feeding, meaning that the fish were fed for a quarter to half the time we used. Elevated levels of anxiety have been associated with increased locomotor activities in zebrafish (MacPhail et al., 2009; Irons et al., 2010; Cachat et al., 2013; Vignet et al., 2013). Thus, HFD-induced anxiety may be accompanied by an increase in locomotion.

## Conclusion

To sum up, our data strongly suggest that consumption of HFD triggers a non-traumatic regenerative response in the zebrafish brain. Since the regenerative capacity of the mammalian brain is very restricted, HFD feeding could be highly detrimental to the mammalian brain. Thus, revealing the molecular mechanisms that initiate the regeneration-like program of the zebrafish brain to HFD exposure may offer a solution to not only fight obesity but also heal non-traumatic injuries of the brain.

## Acknowledgment

We thank Izmir Biomedicine and Genome Center Vivarium-Zebrafish Core Facility and the facility manager Emine Gelinci for providing zebrafish care, Optical Imaging Core Facility and the facility manager Dr. Melek Ucuncu for microscope facility support and Histopathology Core Facility and the facility manager Ece Uzun for her help in ORO staining.

## Statements and Declarations

### Ethics Statement

The animal study protocol was approved by the Animal Experiments Local Ethics Committee of Izmir Biomedicine and Genome Center (IBG-AELEC) (2020-003, 12/02/2020).

### Author Contributions

GO, YA and GC designed the experiments. YA, YKP, OO, DN, and DI performed the experiments. YA, YKP, OO, DN, and GC drafted the manuscript. GO wrote the manuscript. All authors have read and agreed to the published version of the manuscript.

